# Interaction between mutation type and gene pleiotropy drives parallel evolution in the laboratory

**DOI:** 10.1101/2023.01.19.524694

**Authors:** Philip Ruelens, Thomas Wynands, J. Arjan G.M. de Visser

## Abstract

What causes evolution to be repeatable is a fundamental question in evolutionary biology. Pleiotropy, i.e. the effect of an allele on multiple traits, is thought to enhance repeatability by constraining the number of available beneficial mutations. Additionally, pleiotropy may promote repeatability by allowing large fitness benefits of single mutations via adaptive combinations of phenotypic effects. Yet, this latter evolutionary potential may be reaped solely by specific types of mutations able to realize optimal combinations of phenotypic effects while avoiding the costs of pleiotropy. Here, we address the interaction of gene pleiotropy and mutation type on evolutionary repeatability in a meta-analysis of experimental evolution studies with *Escherichia coli*. We hypothesize that single-nucleotide polymorphisms are principally able to yield large fitness benefits by targeting highly pleiotropic genes, whereas indels and structural variants provide smaller benefits and are restricted to genes with lower pleiotropy. By using gene connectivity as proxy for pleiotropy, we show that nondisruptive single-nucleotide polymorphisms (SNPs) in highly pleiotropic genes yield the largest fitness benefits, since they contribute more to parallel evolution, especially in large populations, than inactivating SNPs, indels and structural variants. Our findings underscore the importance of considering genetic architecture together with mutation type for understanding evolutionary repeatability.

## Main text

The distribution of fitness effects of mutations is a prominent determinant of the tempo and mode of adaptation (1). A classic theoretical framework to analyse the distribution of fitness effects of new mutations via their effect on underlying phenotypes is Fisher’s geometric model of adaptation (FGM) (2-4). In FGM, a genotype is depicted as a point in multidimensional Euclidean space, where the axes correspond to traits and the optimal combination of trait values represents maximum fitness in a given environment (**Fig. 1**). Mutations are represented by a vector connecting ancestral and mutant genotype within this phenotypic space. Depending on the magnitude and direction of this vector with respect to the local fitness maximum, the mutant genotype has higher or lower fitness. Mutations are traditionally defined as universally pleiotropic in FGM (4), signifying that mutations may affect all possible trait combinations with equal probability. However, in real organisms, mutations affect only a subset of traits due to their modular organization (2, 5-8). For example, at the protein level, subnetworks of functionally interacting proteins exist (9-11). A commonly used proxy of a gene’s pleiotropy level is therefore its connectivity, i.e. the total number of functional interactions of a gene and its product (12-15). When pleiotropy is universal, the proportion of beneficial mutations and their average effect size both decrease with increasing pleiotropy, resulting in an apparent ‘cost of complexity’ which constrains the rate of adaptation (16). Under modular pleiotropy, these pleiotropic constraints are alleviated, and some empirical and theoretical studies have even suggested that the largest-benefit mutations occur in highly pleiotropic genes (5, 6, 17). This would be the case when changes in multiple phenotypes are required to maximize fitness, which can in principle be realized by a single mutation in a highly pleiotropic gene (**Fig. 1**).

**Figure 1.**
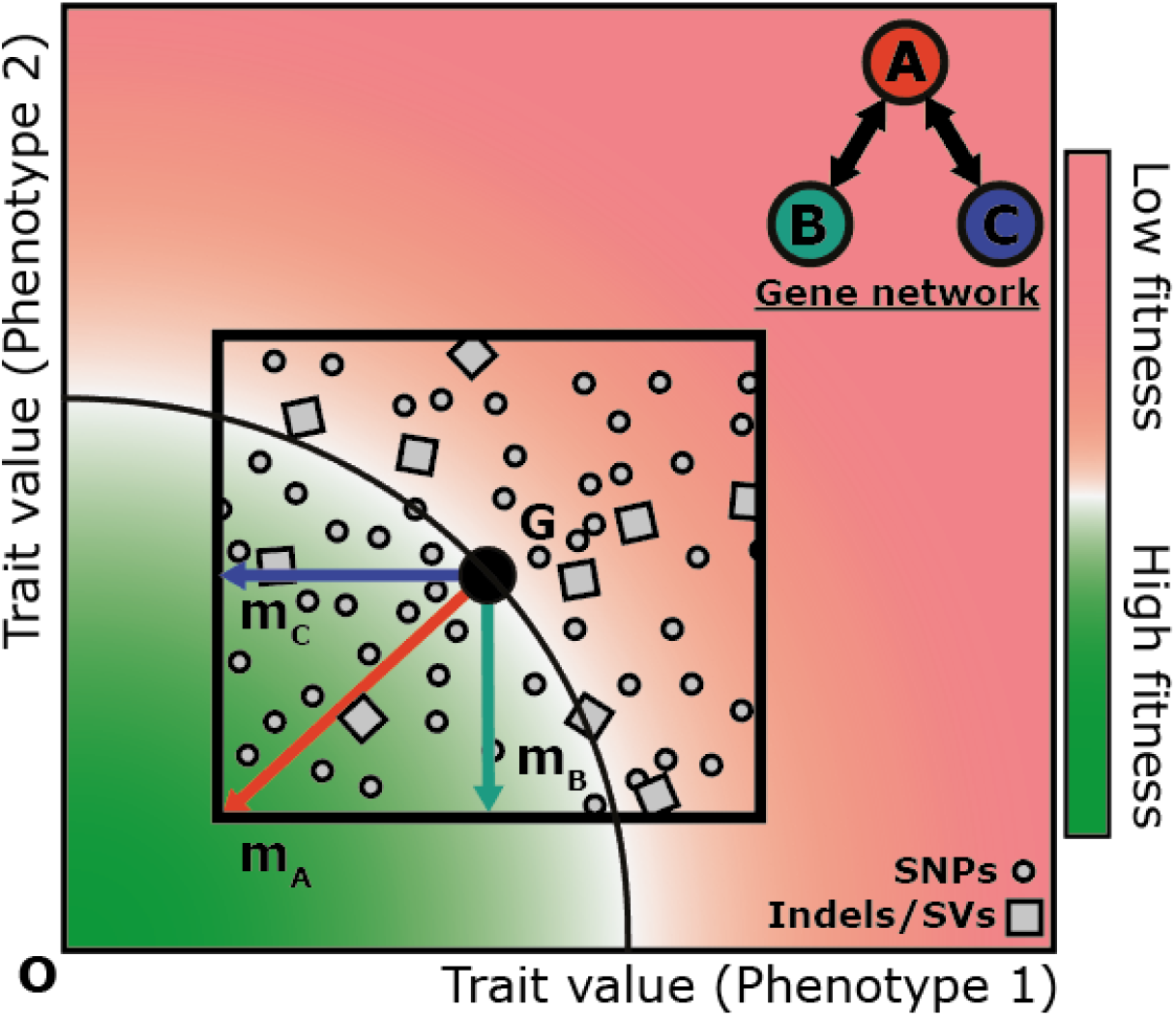
Expected role of gene pleiotropy and mutation type in determining mutational fitness effects under Fisher’s geometric model. Schematic overview of Fisher’s geometric model in two-dimensional trait space with genotype (G) and fitness optimum (O). The fitness of each genotype (shown in color with the black circle sector showing the fitness isocline for G) is determined by the combination of two phenotypes or molecular interactions described by a simplified protein-protein interaction network (top right). Mutations in gene B (with maximum effect m_B_) and gene C (with maximum effect m_C_) affect only one phenotype, while mutations in gene A can theoretically affect both phenotypes and may realize phenotype combinations across the area bounded by the maximum phenotypic effects of alleles in B and C (indicated by the black square). Some beneficial mutations in gene A (with effect m_A_) could optimally combine both single-trait effects and reach higher fitness than mutations in genes B and C alone. Since SNPs (small circles) generate more phenotypically diverse alleles than indels or structural variants (large squares), SNPs sample phenotypic space with greater resolution and are therefore more likely to produce this largest possible benefit. This argument also holds for mutations with larger phenotypic effects that may overshoot the optimum (not shown here), where the greater phenotypic diversity of SNPs allows some to more closely approach the optimum than other mutation classes.

Whether adaptive mutations with large beneficial effects occur more often in genes with high or low pleiotropy remains an open question. Some microbial evolution experiments have found that adaptive mutations with large beneficial effects do occur in highly pleiotropic genes (18-20). However, to realize large fitness benefits in pleiotropic genes, mutations with optimal combinations of multiple phenotypic effects are required. We hypothesize that single-nucleotide polymorphisms (SNPs) are more likely to realize these high-fitness alleles compared to insertions and deletions (Indels) and structural variants (SVs) due to their much greater number and phenotypic diversity compared to other mutation types (**Fig. 1**). Whereas indels and SVs often result in functionally equivalent alleles (loss-of-function or gene amplification), SNPs can fine-tune gene expression as well as functional properties (e.g., catalytic activity, protein stability, DNA-binding, protein-protein interactions). We thus expect SNPs in highly pleiotropic genes to be overrepresented among mutations with the largest benefit that drive parallel adaptation, whereas indels and SVs occur more often in genes with lower pleiotropy where their inherently greater pleiotropic costs are limited.

To test our predictions, we performed a meta-analysis of laboratory evolution studies with the bacterium *Escherichia coli* and examined to what extent gene connectivity, used as a proxy for gene pleiotropy, affected adaptation and parallel evolution of SNPs, indels and SVs. We collected mutational data of 243 independently evolved non-mutator *E. coli* clones from 24 independent evolution experiments, each involving multiple replicate populations (Supplemental data 1 and 2). To allow many different mutation targets to contribute to adaptation, we included only studies using ‘soft selection’ regimes, such as suboptimal growth temperatures (21, 22) and pH levels (23, 24) and novel carbon sources (25, 26). Across all experiments, approximately 27% of *E. coli*’s genes were mutated at least once (1,165 of 4,225 *E. coli* MG1655 genes), thus capturing substantial genomic variation. Finally, given the diminishing role of selection over time (27), we only included studies with a duration between 500-2,000 generations.

To evaluate to what extent gene pleiotropy influenced adaptation, we first determined the connectivity of all target genes in our dataset and compared this to the average connectivity of all *E. coli* genes. We define target genes of each experiment as all mutation-bearing genes with at least one mutation across replicate lines. Estimates of the number of potential physical and functional interactions, or connectivity, of each gene were obtained from the STRING protein-protein interaction database, with a stringent combined confidence score of 0.7 (10). In contrast to expectations from FGM, target genes had a 1.22-fold higher connectivity than the average *E. coli* gene (**Fig. 2A**, Wilcoxon’s test, P<0.0001), suggesting genes with above-average pleiotropy play an important role during adaptation. We also considered for each target gene how often it was mutated, since genes targeted only once may include hitchhiking events of nonadaptive mutations. Genes with multiple independent mutations (≥ 2) per experiment were on average 1.61-fold more connected than genes targeted by just one mutational event (**Fig. 2B**, *t*=-2.86 P=0.009). Moreover, there is a positive relationship between the number of independent mutations within a single gene per experiment and the connectivity of that gene (Generalized linear mixed model with experiment as random factor, *χ*^*2*^=9.78, P=0.002), implying that mutations targeting genes with more connections are more beneficial.

**Figure 2.**
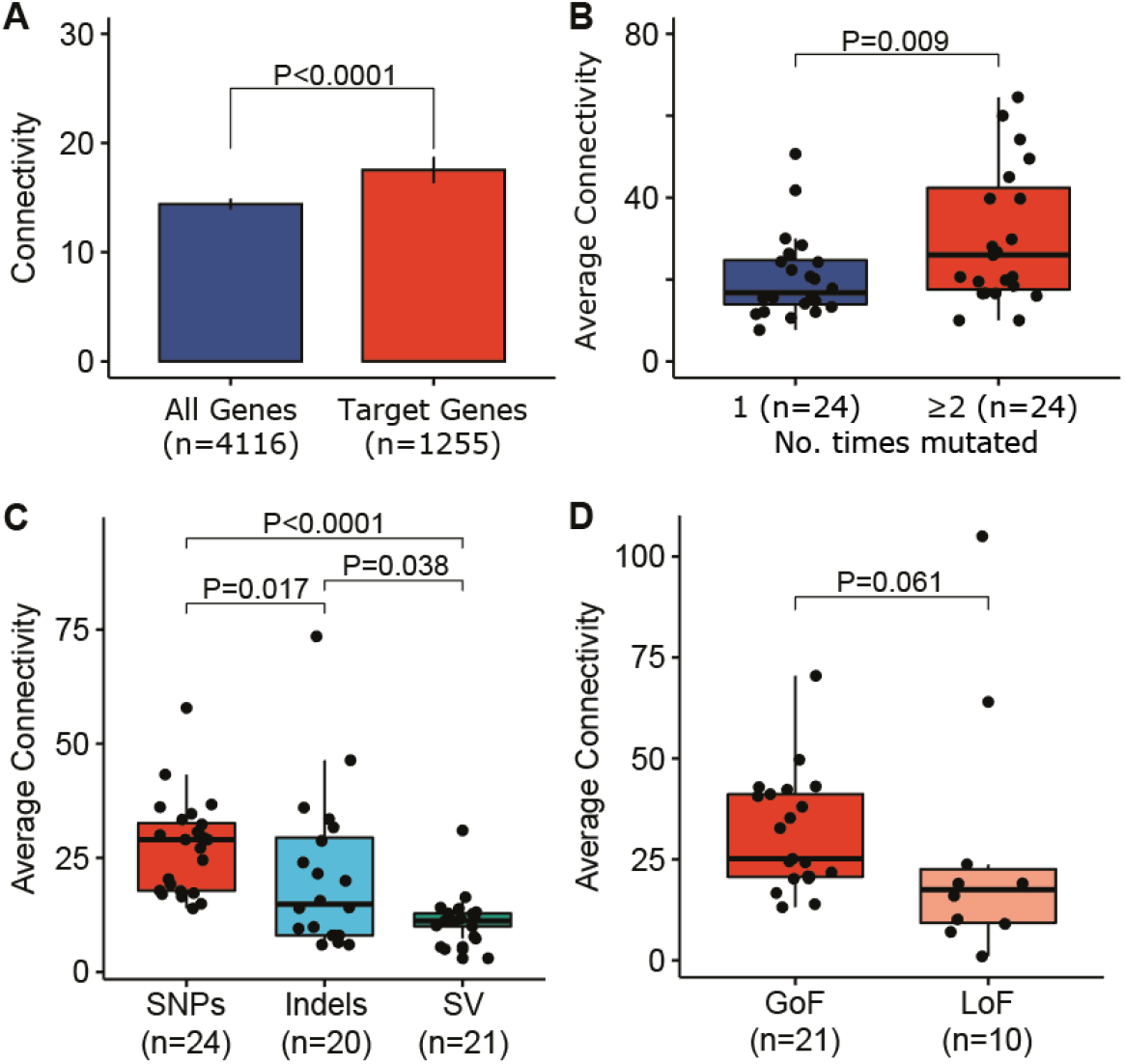
Gene pleiotropy, measured by the level of connectivity, of mutation targets in experimental evolution studies with *E. coli*. **(A)** Connectivity of all *E. coli* genes versus genes targeted by mutations during adaptation. Connectivity estimates were obtained from the STRING interaction database (means ± 95% CI). Statistical significance was determined by a Wilcoxon’s test. (B) Boxplots of average per-experiment connectivity of target genes with one or multiple independent mutations. P-value was determined by a post-hoc contrast test in a mixed-effects model with experiment as random factor. (C-D) Boxplots of the average per-experiment connectivity of genes targeted by different mutation types (C) and by gain-of function (GoF) or loss-of-function (LoF) SNPs (D). P-values are based on post hoc contrast tests (with Tukey’s P-value adjustment) in linear mixed-effects models with experiment as random factor.

Next, we tested the hypothesized interaction between gene pleiotropy and mutation type by determining the average per-experiment connectivity of genes targeted by different mutation types. In total, our dataset consisted of 1,632 mutational events, of which 72% were SNPs, while the remaining mutations were categorized as indels (smaller than 50bp, 20%) and structural variants (8%), which include large genomic rearrangements (>50bp) as well as insertions of insertion-sequence elements. A significant proportion of the variation in the average per-experiment connectivity can be attributed to mutation type (Linear mixed model, *χ*^*2*^=31.17, P<0.0001), with SNP-targeted genes having significantly more interactions compared to SV and indel targets (**Fig. 2C**, post hoc Tukey’s test, *t*=5.57, P<0.0001 and *t*=2.87, P=0.017, respectively), while indel targets were more connected than SV targets (post hoc Tukey’s test, *t*=2.54, P=0.038).

As mutations that inactivate a gene are expected to be less phenotypically diverse than non-disruptive mutations, we asked whether genes affected by putative loss-of-function (LoF) SNPs are less connected than genes affected by gain-of-function (GoF) SNPs. Putative LoF targets were defined as targets where SNPs resulted in a premature stop codon, or where at least one other replicate line within the same experiment showed a frameshift indel or SV, while putative GoF targets were the residual SNP-targeted genes. Although a coarse categorization, we found that the distinction between putative GoF or LoF targets explained a significant amount of variation in target connectivity (**Fig. 2D**, LMM, *χ*^*2*^=4.33, P=0.038), with GoF targets being marginally more connected than LoF targets (t=2.00, P=0.061). We noticed that the LoF category contained two outliers with significantly higher average connectivity compared to the rest of the data. Upon further investigation, we discovered that both these outliers were caused by a possible misclassification within the LoF category of a mutation resulting in a stop codon at position 1367 of the *rpoC* gene. Since *rpoC* has a total length of 1407 amino acids, this might not represent a destructive LoF mutation. Their exclusion from the dataset resulted in a significantly lower connectivity of LoF mutations compared to GoF mutations (t=3.636, P=0.0022). Moreover, it is interesting to note that indels occurred more often in intergenic regions compared to SNPs than expected based on the overall number of mutational events of each type (Chi-square test with Yates correction, *χ*^*2*^=21.9, P<0.0001). Given that indels in protein-coding sequences mostly cause LoF phenotypes, indels may more likely incur beneficial effects when targeting cis-regulatory elements.

Together, these results show that mutations with diverse phenotypic effects, such as (nondisruptive) SNPs, more often than other mutation types fix during adaptive laboratory evolution of *E. coli* when they occur in highly connected genes. Does this mean that SNPs also have larger beneficial effects? To address this, we considered the degree of parallel evolution as proxy for fitness. We analysed the degree of repeatability of fixed mutations from each type by calculating the average pairwise genotype similarity, or *H*-index, at the gene level between independent lines of the same experiment for SNPs, indels and SVs ((28), see supplementary methods). We found a significant effect of mutation type on gene-level repeatability (**Fig 3A**, LMM, *χ*^*2*^=26.73, P<0.0001). Repeatability was significantly higher for SNPs than for both indels and SVs (post hoc Tukey’s test, *t*=4.46, P=0.0002 and *t*=4.49, P=0.0001, respectively), while repeatability of indels and SVs did not differ (post hoc Tukey’s test, *t*=0.033, P=0.99). Among SNPs, whether a mutation target was a putative GoF or LoF target also had a significant effect on repeatability (**Fig. 3B**, LMM, *χ*^*2*^=6.174, P=0.013), with GoF mutations being more repeatable than LoF mutations (*t*=2.23, P=0.044).

**Figure 3.**
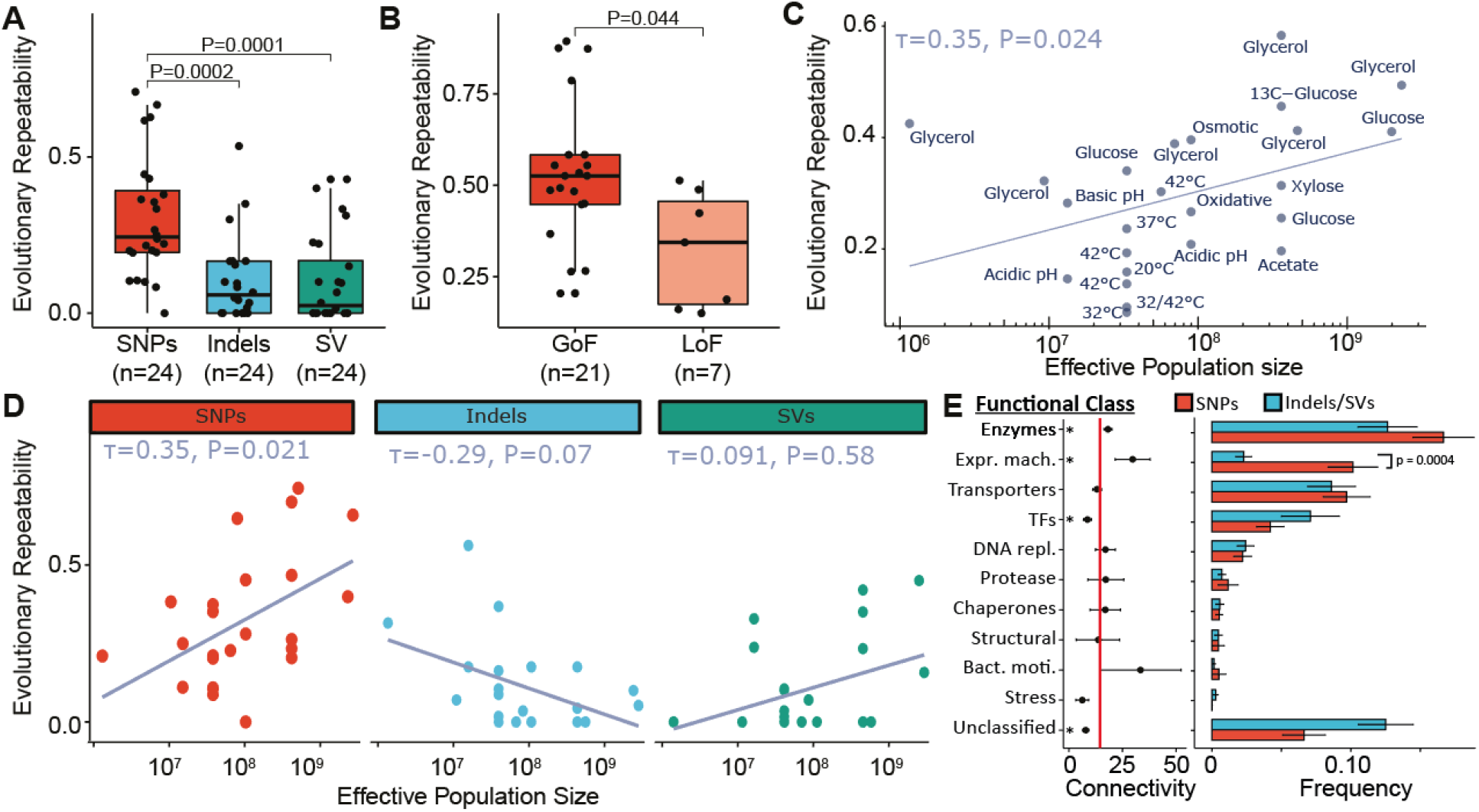
Evolutionary repeatability is driven by large-benefit SNPs, particularly in large populations. Boxplots (A) and (B) show the repeatability at the gene level of different mutation types (A) and GoF and LoF SNPs (B). P-values are based on post hoc constrast tests (with Tukey’s P-value adjustment) in linear mixed-effects models with experiment as random factor. Only significant differences are indicated. (C) Per-experiment evolutionary repeatability at the gene-level depends positively on effective population size, suggesting it is driven by selection rather than mutation bias. Labels show the selective conditions of the different evolution experiments. The Kendall rank correlation coefficient (*τ*) with associated P-value is shown in the top left corner. (D) Dependence of evolutionary repeatability at the gene level of different mutation types on effective population size. The Kendall rank correlation coefficient (*τ*) with associated P-value are shown at the top of each subplot. (E) Per-experiment comparison of SNP and Indels/SV frequencies for genes beloning to different functional classes alongside the average connectivity of each class. Asterisks indicate signficant differences between the connectivity of each functional class and global connectivity of all *E. coli* genes (indicated by the red vertical line). Statistical significance was determined by a Wilcoxon’s test with Bonferroni correction. Abbreviations: Expr. Mach.: Expression machinery, TFs: Transcription Factors, Bact. Moti.: Bacterial motility.

Although natural selection is considered the main driver of parallel evolution, mutation bias (i.e. variation in the rate of different mutations) may also contribute to parallel evolution, 8especially in small populations (28-30). To distinguish between mutation and selection bias as cause of parallel evolution, we examined the dependence of parallel evolution on the population size. We expect a positive dependence if selections dominates parallel evolution, whereas no or a negative effect of population size would indicate an influence of mutation bias on parallel evolution (28). To do so, we estimated the effective population size (*N*_*e*_) of each experiment based on the experimental conditions as *N*_*e*_ ≈ *g * N*_*0*_, with *g* representing the number of generations per serial transfer and *N*_*0*_ the population size transferred (31). Effective population size varied approximately 2,000-fold from 1×10^6^ to 2×10^9^ across studies. Consistent with previous observations and with a more prominent role of clonal interference in large populations (32, 33), evolutionary repeatability increased with effective population size (Kendall’s rank correlation, *τ*=0.35, P=0.024, **Fig 3C**). As the mutation supply and competition between beneficial mutations increases with increasing population size, the efficiency of natural selection to fish out the largest-effect mutations increases, even when they occur at lower mutation rates (28-30, 34). This suggests that selection is the dominant cause of parallel evolution in the included evolution experiments.

When comparing the dependence of evolutionary repeatability on population size for the three mutation types, only the repeatability of SNPs was positively correlated with population size (**Fig. 3D**, Kendall’s rank correlation, *τ*=0.35, P=0.021). This confirmed our expectation that SNPs drive parallel evolution via their greater fitness effects rather than higher mutation rates, compared to indels and SVs. Protein interaction networks are generally scale-free, indicating that their interaction distribution is significantly skewed towards lowly connected genes (35, 36). Despite the more numerous genes with few connections, selection apparently fishes out rare SNPs in higher connected genes, supporting a key role of natural selection driving parallel evolution. These results generalize the findings of a recent evolution experiment with populations of *E. coli* of different size that were challenged with a β-lactam antibiotic (28). There we found that parallel evolution of antibiotic resistance in small populations was mostly driven by SVs, whereas in large bacterial populations SNPs dominated parallel evolution. Inferences of the rates and fitness effects of these mutation classes using simulations to match the experimental results confirmed that the repeatability of SVs and SNPs was driven by high rates and large fitness benefits, respectively. The link with gene pleiotropy reported in the present study also provides a potential explanation for the observed distinct roles of SNPs and SVs in our previous work.

Finally, we explored the function of the genes affected by the different types of mutations. To do so, we clustered all mutated genes in functional classes based on the KEGG BRITE classification system for proteins (37) and subsequently compared the frequency of SNPs and Indels or SVs for genes in each class. This analysis showed that only for genes involved in translation and transcription there was a significant difference between the mutation types, with SNPs being significantly more frequent than indels or SVs in these genes (Fig. 3E, Wilcoxon’s test, P = 0.0004). When comparing the connectivity of different functional classes, only the Expression machinery and Enzyme class showed a significantly increased connectivity compared to the average *E. coli* gene (Wilcoxon’s test, P_adjusted_ < 0.05), while the other classes are either not significantly different or significantly lower (Transcription factors, Stress and Unclassified). Although enzymes were not significantly more targeted by SNPs compared to indels, genes of the expression machinery class were, consistent with our hypothesis that SNPs are best able to realize large fitness benefits in these genes with above average connectivity. The most frequent mutation targets within this functional class are the genes encoding the β and β’ subunits of RNA polymerase, *rpoB* and *rpoC*, respectively. Both genes are highly pleiotropic (38, 39), as they are among the top 1% most connected genes.

Understanding pleiotropy is of profound importance to evolutionary biology, as it is an essential feature of the genotype-phenotype map and may affect both the tempo and mode of evolution (40). Pleiotropy has traditionally been considered as a constraining factor of adaptation (4, 16). However, during laboratory adaptation to novel conditions, selection targeted genes with more functional interactions than average (18-20, 41). We show that this may be caused by the ability of highly connected genes to yield larger beneficial effects, as evidenced by their increased repeatability during early adaptation in a novel environment. Primarily non-disruptive SNPs are able to escape the cost of complexity and capitalize on the greater adaptive potential of highly pleiotropic genes, resulting in large net fitness benefits. By contrast, indels and SVs, due to their limited phenotypic resolution and inability to avoid predominant antagonistic pleiotropic effects, are restricted to genes that affect fewer traits.

It is important to note that the observations and conclusions of this study are based on the adaptation of *E. coli* to stable environments in the laboratory and it is uncertain whether these trends would hold true for other species and under natural conditions. Recent work with Drosophila revealed higher pleiotropy of new spontaneous mutations compared with alleles at higher frequencies that make up the standing genetic variation (42, 43). These observations are still consistent with our conjecture that only the very best alleles that fix – as those in our meta-analysis – are enriched for highly pleiotropic genes, whereas pleiotropic mutations are on average deleterious. However, also differences in effective population size, genetic architecture and selective conditions may play a role. For example, natural populations likely experience more variable conditions, which may counter-select highly pleiotropic mutations with large fitness benefits in a specific environment, as was found in recent studies with *Bacillus subtilis* (44) and yeast (8). These environmental-dependent pleiotropic effects may limit the long-term success of these alleles in fluctuating environments (45). As such, the contribution of large-benefit mutations in highly pleiotropic genes could be limited to adaptation in natural environments, where conditions may fluctuate continuously. Nevertheless, our findings highlight a fundamental interplay between gene pleiotropy and mutation class underlying parallel evolution, which may stimulate further research.

## Methods

### Data collection

We caried out keyword searches (“experimental evolution *Escherichia coli*”) of online scholarly repositories using Google Scholar for studies that performed laboratory evolution experiments with *Escherichia coli* and sequenced evolved clones. As we were interested in the repeatability of genetic changes, included studies needed to have at least two independent non-mutator populations undergoing the same selective conditions for a minimum of 500 generations. Mutator populations were excluded based on their substantially elevated substitution frequencies. Furthermore, mutational data from a single evolved clone per replicate line should have been available as of May 2020. We excluded studies that selected for resistance or tolerance of toxic compounds (antibiotics or other chemical compounds), as we wanted to focus on soft selection conditions where relative fitness and clonal interference are more important in determining evolutionary repeatability than variation in mutation rates (mutation bias). All included studies conducted experimental evolution using serial passage protocols, allowing us to estimate the effective population size as the number of generations per transfer multiplied by the population size transferred (31).

### Gene connectivity

The connectivity of genes was based on the number of high confidence links (combined cut-off >0.7) in the STRING database (v11, *Escherichia coli K12 substr. MG1655 (10)*). The connectivity of intergenic mutations was calculated as the average connectivity of the two neighbouring genes. Genes that could not be mapped to the STRING database were excluded from the connectivity analyses.

### Gene-level repeatability

Repeatability at the gene-level was determined as in ref. (28). Briefly, evolutionary repeatability or H-index is a measure for the fraction of shared mutations of two genotypes, and was calculated by first determining the pairwise H-index of each independently evolved clone versus all other final clones. For the comparison of clones A and B with *m* and *n* mutated genes, the pairwise H-index of A to B is:

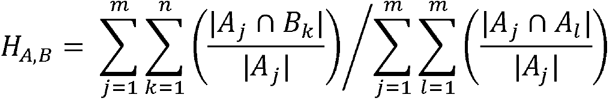

Due to possible asymmetries in the overlap between mutations of different size, such as a SNP and an SV, we calculated also the pairwise H-index of B to A and used the average of both values as the pairwise H-index of clones A and B. The repeatability of an evolution experiment is consequently the mean of all pairwise H-indices among the total set of clones from replicate populations. To determine the repeatability of different mutation types, only genes targeted by the mutation type of interest were considered.

### Functional classification

All genes within our dataset were classified in ten functional classes based on their classification in the upper level of the KEGG BRITE hierarchical organization of ‘Genes and Proteins’ (37). For those genes that could not be mapped to the KEGG BRITE database, their classifications were based on the PANTHER protein classification system or their described function in UniProt (46, 47). Genes that could not unambiguously be grouped in any of the ten functional classes remained ‘Unclassified’.

### Data analysis

Data analysis was conducted using *R* statistical software (v3.6.1). Non-parametric Kendal rank correlation tests were performed using the ‘cor’ function with method ‘kendall’ (base package). Non-parametric comparisons of two unpaired means were evaluated with a Wilcoxon’s test using function ‘wilcox.test’ in R (base package). To assess the relationship between mutation type and average per-experiment connectivity or repeatability, a linear mixed-effects model (LMM) was applied using the ‘glmer’ function (lme4 package, gaussian distribution), where experiment was considered as random factor allowing the intercepts to vary per experiment. A mixed-effect model was preferred over a paired test with P-value adjustment, as the former allows partially unpaired data to contribute to the model’s mean estimates. Log transformation of connectivity and repeatability was performed to transform skewed data to conform to a normal distribution. Post hoc pairwise comparisons of mean estimates were performed using the ‘emmeans’ and ‘pairs’ or ‘contrast’ functions implemented in the ‘emmeans’ package. The relationship between the number of independent mutations (count) and the connectivity of the mutated genes was assessed using a negative binomial generalized linear mixed model (GLMM) using the ‘glmer.nb’ function with the log-transformed count as dependent variable, the log-transformed connectivity as explanatory variable and experiment as random factor (lme4 package). LMM and GLMM assumptions were visually evaluated using the residual diagnostics package: ‘DHARMa’ (48). The data and R script required to reproduce analyses and figures of this study are openly available in the Dryad Digital Repository at https://doi:10.5061/dryad.7h44j0zz4.

## Acknowledgments

We thank Annette Van Oystaeyen for help with statistical analyses. This work was supported by a Human Frontiers in Science Program grant (RGP0010/2015) and a grant from the Netherlands Organization for Scientific Research (OCENW.XS.058) to JAGMdeV and an EMBO fellowship (ALTF 273-2017) to PR.

## Author contributions

Conceptualization, PR and JAGMdeV; Methodology, PR and JAGMdeV; Investigation, PR and TW; Data Curation, PR; Writing – Original Draft, PR; Writing – Review & Editing, JAGMdeV; Funding Acquisition, JAGMdeV and PR.

## Declaration of interests

The authors declare no competing interests.

## Notes

https://doi.org/10.5061/dryad.7h44j0zz4

### Competing Interest Statement

The authors have declared no competing interest.

